# Gene tree discordance causes apparent substitution rate variation

**DOI:** 10.1101/029371

**Authors:** Fábio K. Mendes, Matthew W. Hahn

**Author notes:** Correspondence to be sent to: 1001 E. Third St., Bloomington, IN, 47405, USA.

## Abstract

Substitution rates are known to be variable among genes, chromosomes, species, and lineages due to multifarious biological processes. Here we consider another source of substitution rate variation due to a technical bias associated with gene tree discordance, which has been found to be rampant in genome-wide datasets, often due to incomplete lineage sorting (ILS). This apparent substitution rate variation is caused when substitutions that occur on discordant gene trees are analyzed in the context of a single, fixed species tree. Such substitutions have to be resolved by proposing multiple substitutions on the species tree, and we therefore refer to this phenomenon as “SPILS” (Substitutions Produced by Incomplete Lineage Sorting). We use simulations to demonstrate that SPILS has a larger effect with increasing levels of ILS, and on trees with larger numbers of taxa. Specific branches of the species trees are consistently, but erroneously, inferred to be longer or shorter, and we show that these branches can be predicted based on discordant tree topologies. Moreover, we observe that fixing a species tree topology when performing tests of positive selection increases the false positive rate, particularly for genes whose discordant topologies are most affected by SPILS. Finally, we use data from multiple *Drosophila* species to show that SPILS can be detected in nature. While the effects of SPILS are modest per gene, it has the potential to affect substitution rate variation whenever high levels of ILS are present, particularly in rapid radiations. The problems outlined here have implications for character mapping of any type of trait, and for any biological process that causes discordance. We discuss possible solutions to these problems, and areas in which they are likely to have caused faulty inferences of convergence and accelerated evolution.

---

Characterizing the rate of molecular evolution and the forces driving variation in this rate is a major goal of evolutionary biology. Studying how and why substitution rates vary allows us to make inferences about the biology and evolutionary past of extant species and genes (Bromham 2009). Examples of such inferences include: learning about past episodes of adaptive evolution (e.g., Larracuente et al. 2008; Studer et al. 2008), whether species’ life-history traits influence how fast they evolve (e.g., Martin and Palumbi 1993; Bromham et al. 1996; Smith and Donoghue 2008), and whether specific environments make species prone to higher mutation rates (e.g., Wright et al. 2006). A better understanding of the tempo of molecular evolution can also be used to improve the dating of evolutionary events (Lanfear et al. 2010).

Substitution rates vary at several different scales, including among genes, regions with different recombination rates, chromosomes, species, and clades. Variation among genes can be driven by natural selection. Adaptive evolution increases the rate of substitution, while negative selection decreases it (Li 1997). Even in the absence of differences in selective effects there can be variation in the substitution rate due to variation in the underlying mutation rate. For instance, regions of low recombination have been found to have lower substitution rates (Hellmann et al. 2003). This association has been explained by the mutagenic effects of recombination (Hellmann et al. 2003) or by linked selection in recent ancestral populations (Begun et al. 2007). Substitution rates also vary among chromosomes, with sex chromosomes such as the X chromosome in mammals exhibiting lower substitution rates (e.g., Shimmin et al. 1993; Chang et al. 1994; Ebersberger et al. 2002; Malcom et al. 2003; The Chimpanzee Sequencing and Analysis Consortium 2005). Male mammals undergo more germline cell divisions than females, and therefore because the X spends less time in males it accumulates fewer mutations (Miyata et al. 1987; Makova and Li 2002). Species also vary in their rate of substitution, for which several underlying forces have been put forward. Possible causes include variation in the metabolic rate (Martin and Palumbi 1993), generation time (Laird et al. 1969; Wu and Li 1985), population size (Lynch 2010), and body size (Bromham et al. 1996). Finally, the number of speciation events in a clade is correlated with its substitution rate, leading to the assertion that speciation is an active force promoting bouts of accelerated evolution (Webster et al. 2003; Pagel et al. 2006; Venditti and Pagel 2009; Zhang et al. 2014).

Here we consider another source of apparent substitution rate variation due to a technical bias associated with gene tree discordance. Gene trees can differ in topology from the species tree because of diverse phenomena such as incomplete lineage sorting (ILS), hybridization, and lateral gene transfer (Maddison 1997). Gene tree discordance – differences between gene trees and the species tree – caused by ILS is common in nature (Pollard et al. 2006; Scally et al. 2012; Brawand et al. 2014; Jarvis et al. 2014; Zhang et al. 2014; Suh et al. 2015), with ILS levels being proportional to ancestral population sizes and inversely proportional to time between speciation events (Pamilo and Nei 1988; Maddison 1997). ILS can occur among species at any point in the past, and therefore can affect phylogenetic inference across all timescales (Oliver 2013).

How does gene tree discordance lead to apparent substitution rate variation? If a single species tree is used for analysis, nucleotide substitutions that occur on internal branches of discordant gene trees that are absent from the species tree must then be placed on existing species tree branches (Fig. 1). All bifurcating discordant gene trees contain one or more internal branches that are not present in the bifurcating species tree (see e.g. Fig. 1; Robinson and Foulds 1981). Substitutions occurring on these internal branches of discordant gene trees (filled symbols in Fig. 1) must be accounted for by multiple substitutions on the species tree (hollow symbols in Fig. 1). Consider the filled cross (representing a substitution from 0➔1) on discordant tree 1 (Fig. 1). Because taxa A and C now both have state 1, on the species tree this substitution must be explained by multiple substitutions, in this case an initial 0➔1 change on the branch subtending A, B, and C (hollow cross in Fig. 1) and a back substitution on branch B (also represented by a hollow cross in Fig. 1). The same reasoning applies to substitutions occurring on internal branches of discordant gene tree 2 that do not exist in the species tree (circles in Fig. 1). No matter how many taxa are involved, when substitutions on discordant gene trees are analyzed in the context of a single species tree, spurious substitutions will be inferred. Therefore, with more discordant gene trees included in an analysis (i.e. the more ILS is present), we expect a higher inferred substitution rate.

**Figure 1.**
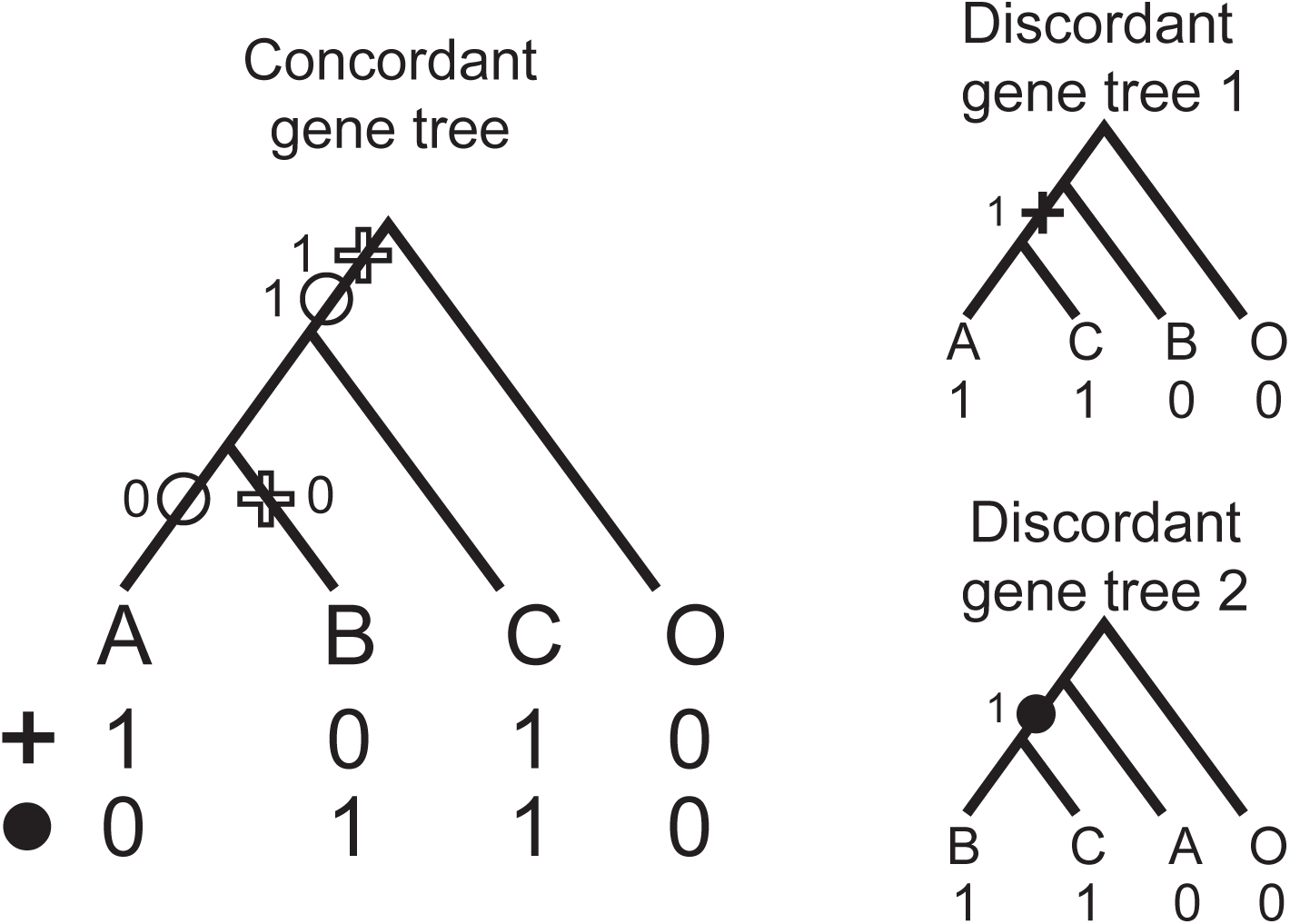
Species tree and two discordant gene trees. The filled cross and filled circle on gene tree 1 and 2, respectively, correspond to 0➔1 substitutions that occurred in two different hypothetical loci. When both genes are analyzed using the species tree, these substitutions are incorrectly mapped (hollow symbols) on the branch leading to ((A,B),C) and as reversals on either branch A or B. As a result, branches A, B, and the branch leading to ((A,B),C) get spuriously longer.

The technical bias described above has been previously considered in a wider context, irrespective of the character’s nature, and dubbed “hemiplasy” (Avise and Robinson 2008). The rationale behind this term is that a homoplasy-like effect is observed (i.e., character transitions are mapped to two or more branches mimicking convergent evolution or character-state reversal), yet the event is not actually homoplasious (Avise and Robinson 2008). In the particular case of analyses of sequence data in the presence of ILS, we hypothesized that there would be consistent biases and predictable patterns of substitution rate variation. To distinguish this phenomenon from the general pattern of hemiplasy we henceforth refer to it as SPILS (Substitutions Produced by Incomplete Lineage Sorting), returning later to discuss the relationship between the two. In the present work we use simulated data and data from multiple *Drosophila* species to demonstrate the magnitude and ubiquity of SPILS. We indeed observe predictable biases, including consistent patterns of apparent substitution rate acceleration and deceleration on individual branches, and an effect of SPILS on increasing the false positive rate in tests for positive selection.

## Materials and Methods

### Quantifying the Effects of SPILS on Branch Lengths

In order to test whether ILS leads to inferences of spurious substitutions, we simulated 1-kb sequences using a population size of *N*_e_=10^6^ and a mutation rate of *μ*=10^−9^. There was no recombination within loci. Ten thousand (10,000) sequences were simulated along species tree (((A,B),C),D) and ((((A,B),C),D),E) under 4 different ILS conditions, named ILS0, ILS1, ILS2 and ILS3, in order of increasing gene tree discordance starting from no ILS (ILS0). The amount of ILS was increased by changing the *N*_e_ of the internal branches of the two species trees (Supplementary Fig. 1): in the first, three taxa are involved in the discordance [((A,B),C)], and three different topologies are possible (Supplementary Fig. 2a). In the second, four species are involved in the discordance [(((A,B),C),D)], and fifteen different topologies are possible (numbered according to their probabilities; Supplementary Fig. 2b). Simulations were done using the *egglib* package (Mita and Siol 2012).

**Figure 2.**
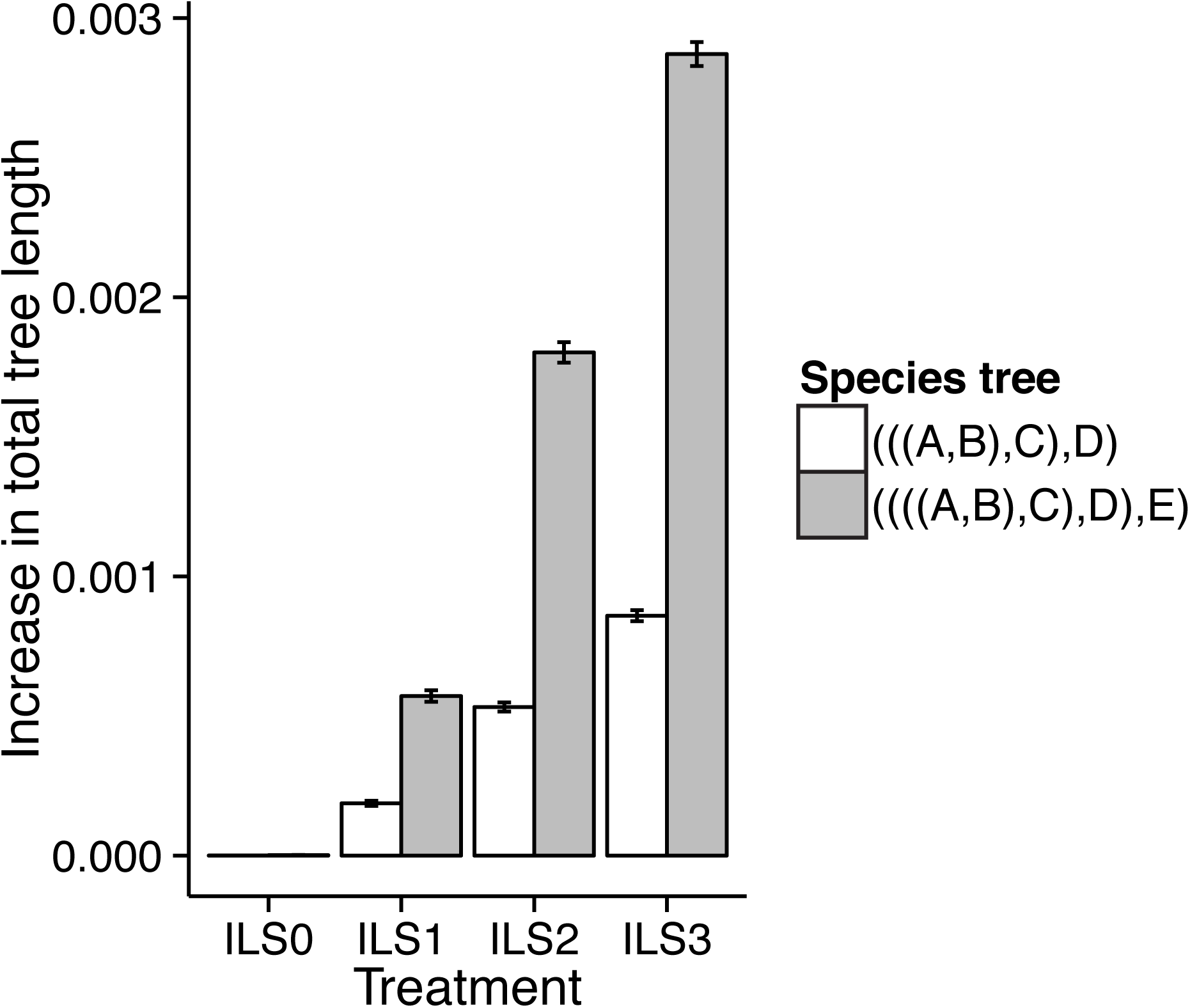
Mean total tree length increase (calculated as the difference in length of “Fixed” topologies and “Unfixed” topologies) by ILS treatment, for ILS among ((A,B),C) and (((A,B),C),D). Based on 10,000 simulated topologies for each treatment.

For each simulated alignment, branch lengths were inferred using two kinds of trees, fixed (F) and unfixed (U), using PhyML (Guindon and Gascuel 2003) with the GTR nucleotide sequence model. “F” trees were constrained to be topologically equivalent to the species tree, that is, they were forced to be either (((A,B),C),D) or ((((A,B),C),D),E). “U” trees were free to have the topology that maximized their likelihood given the alignment.

We quantified the effects of SPILS by comparing the lengths of F and U trees, as SPILS should make the former spuriously longer. Comparisons are made between these two conditions rather than with the known, simulated topology because with increasingly more ILS, terminal branches tend to become longer and internal branches shorter as a result of deeper coalescence in the ancestral populations (Gillespie and Langley 1979). These length differences are not caused by SPILS, and are not the subject of this study (see Angelis and dos Reis 2015 for a discussion of this problem).

In order to detect an increase in total tree length due to SPILS, we calculated total tree length differences (F – U) for each of the 10,000 simulated alignments. We then proceeded to determine which branches were affected using two approaches. First, for each F tree branch, we compared its average length across all 10,000 sequences in the ILS3 condition for the ((A,B),C) and (((A,B),C),D) scenarios. Second, again for each F tree branch, we looked for any increase in length by calculating (F – U) branch length differences for all 10,000 sequences, and then grouped these differences by topology.

Both these approaches must deal with the complication that discordant trees differ from the species tree by one or more internal branches that do not exist in the species tree, making these branches hard to compare. Given species tree ((A,B),C), for instance, an internal branch subtends (A,B); but in the discordant topologies resulting from ILS the internal branch subtends either (A,C) or (B,C). It is precisely these internal branches lacking a counterpart in the species tree that give rise to SPILS – substitutions on these branches cannot be directly mapped to the species tree’s internal branch, which necessitates mapping to multiple other branches. Therefore, we employed the same procedure used for terminal branches (calculation of F – U differences), but with a key difference: when allowing genes to take their own trees (U trees), we used the lengths of the internal branches present in those trees (and absent in the species tree).

### Quantifying the Effect of SPILS on Inferences of Positive Selection

The three and fifteen different topologies resulting from ILS among ((A,B),C) and (((A,B),C),D), respectively, were used to simulate 999-bp coding sequences with *evolver*, a program from the PAML suite (Yang 2007). One thousand coding sequences were simulated for each topology, with 60% and 40% of the sites being constrained (ω = 0.1) and unconstrained (ω = 1), respectively. No sites were simulated with positive selection.

Branch lengths used in the *evolver* simulations were the average lengths obtained from the ILS3 scenario when either ((A,B),C) or (((A,B),C),D) were the clades experiencing ILS. Positive selection was inferred with M1a-M2a site tests (as implemented in PAML v4.7a, [Yang 2007]), specifying species tree (((A,B),C),D) or ((((A,B),C),D),E) as the topology to use regardless of the topology employed in the simulations. We counted a gene as under positive selection using a likelihood ratio test between model M1a and M2a, with 2 degrees of freedom and *P*<0.05 assuming a χ^2^ distribution.

### Quantifying the Effects of SPILS on Drosophila

8,565 gene sequence alignments from the four *Drosophila* species, *D. yakuba, D. erecta, D. melanogaster,* and *D. ananassae* (henceforth abbreviated Y, E, M, and A) used in Larracuente et al. (2008) were obtained from ftp://ftp.flybase.net/genomes/12_species_analysis/clark_eisen/alignments/melangaster_group.guide_tree.longest.cds.masked.tar.gz. Each alignment was used to produce two gene trees each, similar to the simulations described above: F trees had their topology fixed to the species tree topology [(((Y,E),M),A)], and U trees were free to take the topology that maximized the gene tree likelihood in PhyML.

In order to investigate how the presence of intragenic recombination in the *Drosophila* data could influence the effect of SPILS, we concatenated segments of alignments simulated with *evolver* under the three *Drosophila* alternative topologies into a total of 3,000 999-bp “mixed” alignments. Branch lengths used in these simulations were the averages across all *Drosophila* genes trees of each specific topology. These mixed alignments had 70% of their sites from one of the three alternative topologies, and 15% from each of the other two topologies.

## Results

### Total and Individual Branch Lengths

We generated 10,000 simulated alignments for each of the eight ILS scenarios: ILS0, ILS1, ILS2, and ILS3 with either ((A,B),C) or (((A,B),C),D) under ILS. To quantify the effects of SPILS, each simulated alignment was used to infer a pair of trees, either requiring the topology to be fixed (F) to the species tree, or allowing it to be freely inferred (unfixed, or U). Each scenario differed with respect to the level of gene tree discordance exhibited. ILS0, ILS1, ILS2 and ILS3 produced increasingly larger numbers of discordant gene trees, with approximately 0%, 10%, 30%, and 45% of discordant gene trees in each condition when ((A,B),C) was allowed to vary, and 0%, 20%, 50% and 70% when (((A,B),C),D) was allowed to vary (Supplementary Fig. 2). As increasing the number of species allowed for more possible discordant topologies, we predicted it would also lead to more SPILS.

The mean change in total tree length due to SPILS is shown in Figure 2. If SPILS occurs, F trees should be longer than U trees for the same alignment, and this is exactly what we see. The mean change in total tree length (F - U) was positive, and became increasingly larger with increasing levels of discordance. The effect of SPILS was greater when more taxa were involved in the discordance, as trees were inferred to be disproportionately longer when four taxa were involved in the ILS compared to cases where only three taxa were involved (Fig. 2).

Specific branches are expected to become longer or shorter due to SPILS. SPILS is largely caused because some internal branches exist on discordant trees that do not exist on the species tree, whereas some internal branches on the species tree do not exist on the discordant trees. Substitutions that occur on discordant-only branches are mapped twice onto the species tree, causing certain branches to lengthen. While the details of this mapping are surely dependent on the models of sequence evolution used and on the length of species tree branches, we have found that certain branches consistently get longer: PhyML favored a substitution on the ancestral branch subtending the clade undergoing ILS, followed by a reversion on an external branch in all cases (similar results were found with RAxML, [Stamatakis 2014]; results not shown). For example, assuming the species tree ((A,B),C), analyzing a gene that has the discordant gene tree ((A,C),B) should cause branch B and the branch leading to ((A,B),C) to become longer due to SPILS (Fig. 1). If the discordant gene tree ((B,C),A) is used, then branch A and the same branch subtending ((A,B),C) should get longer (Fig. 1). Conversely, because the internal branch subtending (A,B) in the species tree does not exist in either of the two discordant topologies for a rooted three-taxon tree, no substitutions can occur on this branch in these trees and we expect it to therefore become shorter (Fig. 1).

Comparing the average F and U branch lengths from the three and fifteen alternative topologies allowed us to confirm our predictions for branch length changes in the species tree due to SPILS. In the case where only three topologies are possible, branches A, B, and that leading to ((A,B),C) became slightly longer as expected, and the internal branch subtending (A,B) became slightly shorter (Fig. 3a, left panel). Length changes are more pronounced when fifteen alternative topologies are possible (Fig. 3a, right panel), possibly because a greater proportion of the total tree length can be involved in the discordance (see below).

**Figure 3.**
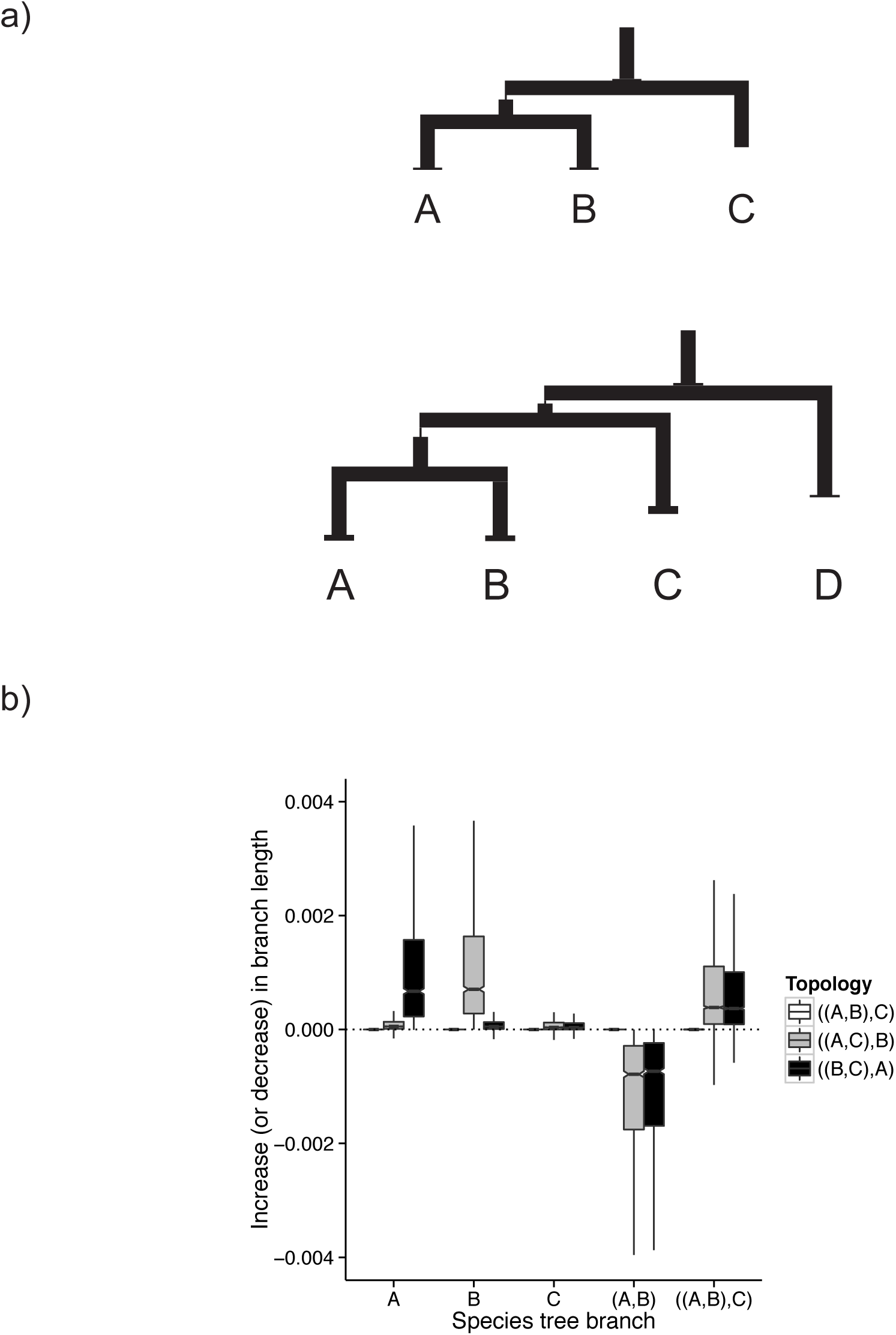
Branch-specific effects of SPILS using simulated data. a) Average species tree branch lengths for ILS among ((A,B),C) and (((A,B),C),D). Wider segments represent the increase in length due to SPILS, while thinner lines represent the decrease in length due to SPILS. b) Increase or decrease in branch length by topology for ILS among ((A,B),C).

Breaking down the effect of SPILS by topology allows us to further explore which branches are affected given alternative gene trees. As predicted, when the discordant gene trees were ((A,C),B) or ((B,C),A), branches A and B became longer due to SPILS, respectively (Fig. 3b). Moreover both topologies showed the ancestor of A, B, and C becoming longer due to SPILS (Fig. 3b). Also as predicted, the number of substitutions by which the internal branch subtending (A,B) gets shorter matches the length increase of the branch subtending ((A,B),C) and either A or B (Fig. 3b). Supplementary Figure 3 shows the predictions for the case of fifteen alternative topologies and Supplementary Figure 4 their confirmation.

**Figure 4.**
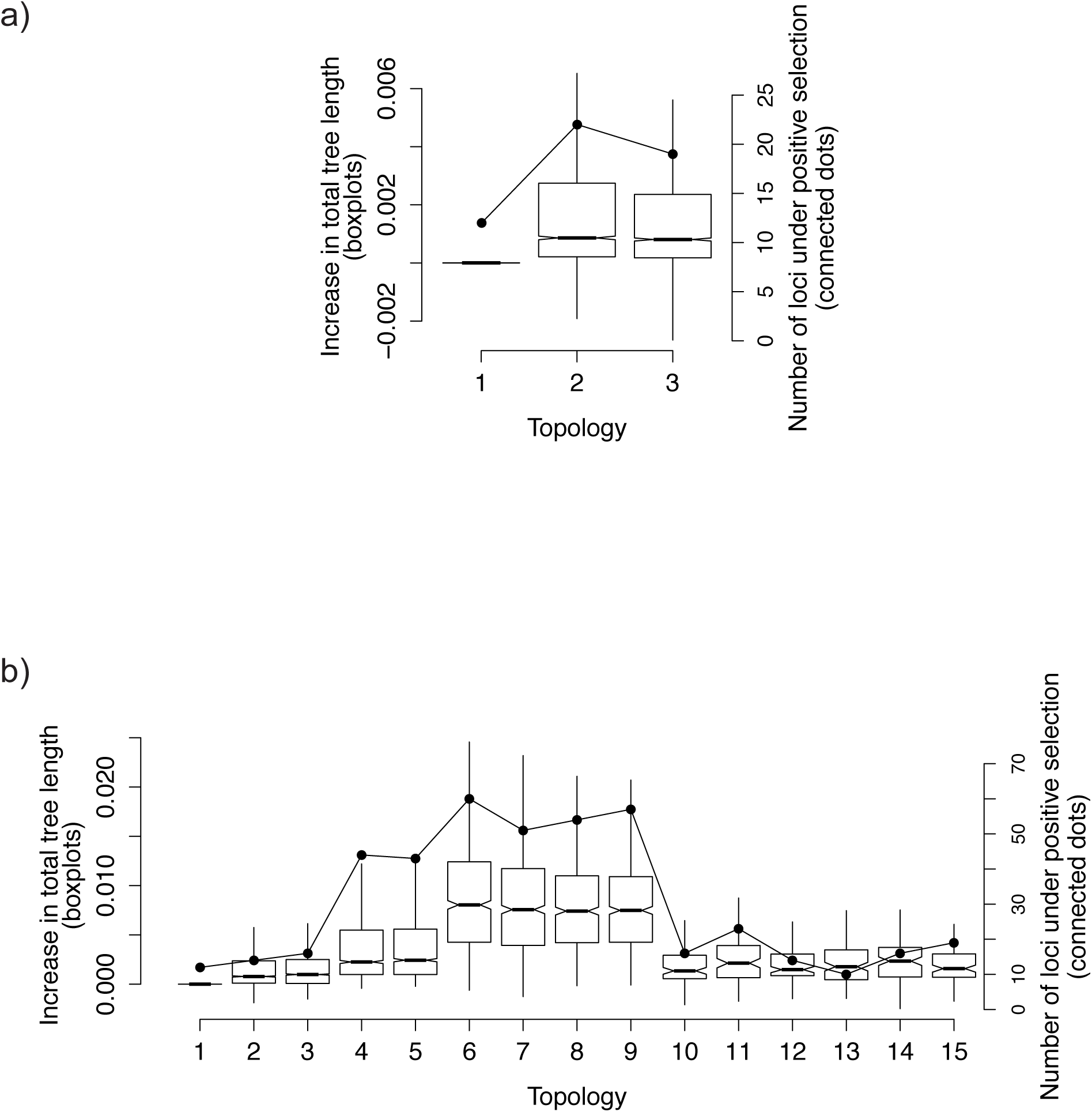
Total tree length increase or decrease (F – U) by tree topology (left axis). Number of genes inferred to be under positive selection using codon site models and a likelihood ratio test (right axis). a) ILS among ((A,B),C). b) ILS among (((A,B),C),D).

Finally, the overall effect of SPILS varied among trees with different discordant topologies, though this was only apparent for the case where ILS involved four taxa. Discordant topologies with a greater number of internal branches absent from the species tree, and with a greater total length of internal branches absent from the species tree, were more affected by SPILS (Fig. 4). Both these gene tree characteristics underlie the effect of SPILS because in fixed-tree analyses longer internal branches absent in the species tree will cause a larger number of substitutions to be misplaced on other branches, causing them to lengthen (Supplementary Fig. 5).

### Inferences of Positive Selection

The results presented above demonstrate that using a fixed species tree when analyzing a collection of possibly discordant gene trees leads to incorrect inferences about the number of substitutions that have occurred. We reasoned that such a phenomenon might also affect inferences of positive selection when studying protein-coding genes. Though nonsynonymous substitutions (as measured by *d*_N_) and synonymous substitutions (*d*_S_) are expected to be affected equally by SPILS, the increased variance in number of substitutions of each type may result in some genes accumulating multiple excess nonsynonymous substitutions. This could in turn lead to spurious inferences of positive selection on individual genes.

To test this idea we simulated protein-coding sequences along topologies that arise from ILS within the species tree ((A,B),C) and (((A,B),C),D) (Methods). To detect positive selection we used the “site” models implemented in PAML (Yang 2007), fixing the tree to be analyzed as the species tree in each case. The genes experienced only negative selection in the simulations, and we therefore expect no positive selection to be detected (or rather, we should only find as much as expected under our null hypothesis of α=0.05).

Overall, we found an excess of genes inferred to be under positive selection when discordant trees were analyzed using the species tree. When there is no discordance the fraction of genes under positive selection was 1.2% for both the three-taxon and four-taxon rooted trees. However, when all topologies are included, the fractions of genes under positive selection were 1.7% and 3% for the same two species trees. Topologies more affected by SPILS (for the reasons discussed above) were particularly more prone to be erroneously classified as being under positive selection: we found a close association between the total increase in tree length due to SPILS and the fraction of trees incorrectly inferred to be under positive selection (Fig. 4b). The specific discordant topologies most affected by SPILS (topologies 6 – 9) showed false positive rates >5% (Fig. 4b). The concordant topology also showed a very slightly lower value of estimated *d_N_/d_S_* than the discordant topologies (0.491±0.005 *vs.* 0.496±0.001).

Interestingly, the topologies that are most affected by SPILS – and also most responsible for the higher false positive rate – are not necessarily the most (or least) frequent of all the possible topologies in the presence of ILS. Topologies numbered 10 to 15 are the five least frequent topologies among the 15 possible ones when (((A,B),C),D) undergoes ILS (Supplementary Fig. 2b), but the number of false positives from these topologies was similar to that from the most frequent discordant topologies (numbered 2 and 3 in Fig. 4b). These results indicate that although the overall influence of SPILS on inferences of positive selection is modest, for certain topologies it may have a much larger effect.

### Drosophila Data

We estimated branch lengths for 8,565 genes from three *Drosophila* species (and an outgroup) known to exhibit considerable levels of ILS (Pollard et al. 2006). We followed the same procedure used for the simulated data, either fixing the tree topology to (((Y,E),M),A), or allowing each gene to take on its own topology. Note, however, that in this case we do not know whether the inferred trees for each gene are the correct topologies; this fact becomes important below.

Our specific predictions are: that total tree lengths from trees fixed for the species tree topology will be longer for genes with discordant trees, that terminal branches Y and E, and the branch subtending ((Y,E),M), will get longer, and that the internal branch leading to (Y,E) will get shorter. Our predictions were confirmed in the *Drosophila* data (Fig. 5a-b). When analyzed under a fixed species tree topology, total tree lengths of genes with discordant gene trees ((Y,M),E) and ((E,M),Y) are inferred to be 2.1% and 1% longer than when each gene is allowed to take on its most likely topology. We also see that the terminal branches leading to Y and E, and the internal branch leading to the common ancestor of ((Y,E),M), are all longer in the fixed-tree analysis, while the internal branch leading to the (Y,E) ancestor is shorter (Fig. 5b).

**Figure 5.**
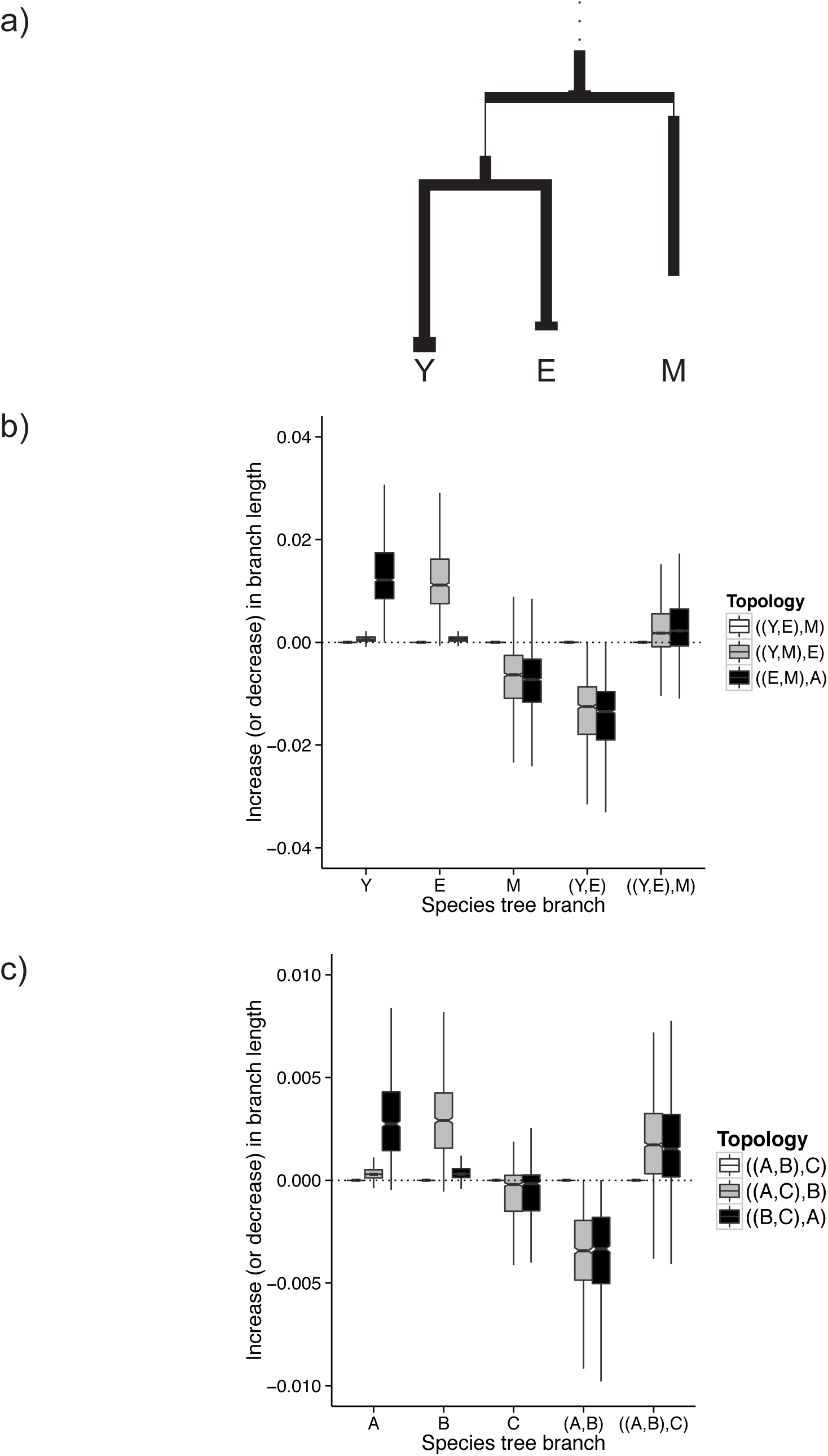
Branch-specific effects of SPILS using *Drosophila* data. a) Average species tree branch lengths for ILS among ((Y,E),M). b) Increase or decrease in branch length by topology for *Drosophila* sequences. c) Increase or decrease in branch length by topology for “recombinant” simulated sequences.

We noticed two differences in these data compared to the simulated data above. First, it often appeared that the external branch leading to species M was shorter in the fixed-tree condition (Fig. 5a), and that this was due to discordant gene trees (Fig. 5b). Second, the increase in length of the branch leading to the ((Y,E),M) common ancestor was smaller than length increases in the Y and E branches (Fig. 5a, b), though these were of similar magnitude in our simulated datasets (Fig. 3b).

We believe that intragenic recombination is the reason for the difference between the results from simulated data and from the *Drosophila* data. SPILS results from assuming one tree when analyzing genes whose sites have for the most part evolved along different topologies as a result of ILS. Recombination within genes, however, means that most genes will have a mix of sites from different topologies; regardless of the major topology inferred for each gene, it is likely that a minority of sites have evolved along another topology. Therefore, even in our “unfixed” trees, enforcing a single (possibly discordant) topology for each gene means that some fraction of sites will show a SPILS-like pattern. The fact that we are now enforcing a different topology means that a different set of branches may be inferred to be longer or shorter. In fact, we can recapitulate the patterns seen in the *Drosophila* data by generating simulated data with intragenic recombination. We generated “recombinant” sequences where 70%, 15%, and 15% of the sites within loci simulated along the three possible topologies using branch length averages obtained from *Drosophila* gene trees. Performing the same comparisons as before recovers the same qualitative and quantitative patterns of branch length differences as in the *Drosophila* genes (Fig. 5c).

## Discussion

The increasing availability of genomic data from multiple species has revealed substantial variation in nucleotide substitution rates (Bromham 2009; Lanfear et al. 2010). It is now recognized that there are a multitude of factors potentially contribution to variation in substitution rates, including variation in functional importance among genes, sex-related differences in mutation rates among chromosomes, variation in life-history traits among species, and even variation in the number of speciation events among clades.

In the present study we systematically characterized how a phenomenon we refer to as “SPILS” affects substitution rates. This phenomenon is expected to occur in any analysis where a fixed species tree topology is used to analyze rates of substitution, but where gene tree discordance is present among the genes being analyzed. SPILS occurs because substitutions on internal branches of discordant gene trees are mapped to multiple species tree branches when the topology is fixed, artificially inflating the length of these branches but shortening the length of species tree branches that are not present in discordant gene trees. Although the effects of SPILS are modest per gene, with more genes, more species, and more discordance present in any dataset, the larger the effects are expected to be. Additionally, we demonstrate that the branches that become spuriously longer (and shorter) due to SPILS can be predicted depending on the discordant gene tree topologies.

Interestingly, the two factors that most affect the degree of discordance due to ILS – the time between successive speciation events and population size – are also often observed to co-vary with substitution rates. SPILS may therefore contribute to variation in substitution rates at many levels. For example, genes located in genomic regions with lower recombination rates may have lower rates of substitution (Hellmann et al. 2003; Begun et al. 2007). Due to an increased effect of linked selection in these regions there is also a lower effective population size and less discordance due to ILS (Pease and Hahn 2013), and therefore less of an effect of SPILS. Similarly, the X chromosome has a lower effective population than the autosomes, and at least in mammals also has a lower substitution rate (Makova and Li 2002; Wilson Sayres et al. 2011). Many species differ in effective population size, and again larger sizes are associated with higher rates of nucleotide substitution (Lanfear et al. 2014). Though there are multiple life-history traits that are correlated with population size (Bromham 2009), we hypothesize that population size itself may drive a positive correlation with substitution rates via SPILS. Finally, clades containing a larger number of species will by necessity have more, shorter internal branches between speciation events, which can result in more ILS and gene tree discordance. It follows that, in addition to any intrinsic effect speciation may have on rates of molecular evolution (Pagel et al. 2006), SPILS is expected to increase the substitution rate of clades with more species.

Although SPILS is expected to occur and to affect substitution rates in all of the aforementioned cases, it does not explain all the variation in any of them. We know this because it is possible to study substitution rates without any effects of SPILS. When only analyzing pairs of species (with or without an outgroup) gene tree discordance is not possible, making SPILS impossible. This analysis has also been suggested in order to control for the node density effect when measuring substitution rates (Witt and Brumfield 2004; Hugall and Lee 2007). Pairwise comparisons have been used to calculate substitution rates among regions differing in recombination rate (e.g., Hellmann et al. 2003), among sex chromosomes and autosomes (e.g., The Chimpanzee Sequencing and Analysis Consortium 2005), among species with different population sizes (e.g., Wu and Li 1985), and among clades with different numbers of species (Barraclough and Savolainen 2001; Lanfear et al. 2010). In each case the expected effect was observed; therefore SPILS cannot explain the entire effect, though it may change the size of the effect observed. However, in cases in which SPILS could affect inferred substitution rates or branch lengths, using only pairs of species – or other possible species trees where no discordance is possible – will ensure no effect of SPILS. In fact, we were able to recover the expected branch lengths from our simulations when removing all but two species and the outgroup (results not shown).

Another strategy for eliminating or minimizing the effect of SPILS is to make an individual tree for each gene to be analyzed. Such an approach is especially relevant when testing each individual gene for positive selection, as we have demonstrated that SPILS causes an increase in false positives. While to our knowledge this problem had not been explicitly discussed before, it is clear that the problem was implicitly recognized in multiple studies. For instance, Larracuente et al. (2008) and Good et al. (2013) inferred trees for each individual gene before testing for positive selection, whereas Scally et al. (2012) simply masked discordant sites consistent with ILS. We note, however, that the former strategy is not optimal because it mitigates the effect of SPILS but does not completely remove them. Within-gene recombination means that any gene likely contains sites that have evolved across multiple different topologies. We observed exactly this pattern in our analysis of the *Drosophila* data, though we have not quantified its effects on inferences of positive selection.

There are additional problems with the strategy of making a tree for every gene. First, in some cases where SPILS is of great magnitude, there may not be enough informative substitutions to accurately reconstruct the history of individual genes. One important instance in which this likely occurs is during species radiations. Radiations involve multiple, closely spaced speciation events, and as a consequence discordance may involve many taxa (e.g., Brawand et al. 2014; Lamichhaney et al. 2015; Suh et al. 2015). There will likely be very few genes that contain substitutions that are informative about all speciation events during a radiation. Furthermore, radiations are expected to be especially hard hit by SPILS because of high levels of discordance among a large number of species, with an outsized accumulation of substitutions on the branch subtending the radiation. In fact, just such a pattern was observed in the radiation of African cichlids (Brawand et al. 2014), with an apparent acceleration of both nucleotide substitutions and gene duplications on this branch. Our results suggest that this pattern may be in part an artifact caused by SPILS.

The strategy of generating individual gene trees may not even be applicable to all situations that are affected by SPILS. Any method attempting to assess the rate of substitution across a tree, for any kind of molecular change, should be affected by SPILS. For example, studies of the rate of gene gain and loss (e.g., Han et al. 2013) or the rate of karyotype evolution (e.g., Jónsson et al. 2014) could both be affected by SPILS, but it is unclear what individual trees one could make to account for discordance. There is no clear single locus or region that all such changes should track, and it may be difficult or impossible to find a region associated with each individual change to make a tree from. In these cases – and in the case of radiations – we suggest that reducing the number of taxa being studied in order to eliminate discordance would lead to more accurate inferences, even though some data is being excluded. Certainly such an approach would make it possible to test for an accelerated rate of evolution on branches leading to radiations, with little loss of power.

As mentioned in the Introduction, SPILS is just one manifestation of the larger problem of hemiplasy (Avise and Robinson 2008). Character-mapping in the presence of ILS will result in incorrect inferences for any kind of character, molecular or not (Hahn and Nakhleh in prep.). These inferences often involve the appearance of convergent changes when none has actually occurred, and we predict that there will be many individual cases of molecular convergence that are ascribable to ILS (Parker et al. 2013). While the problem of mapping character changes onto species trees with underlying discordance has generally gone unaddressed, interestingly it has been addressed in multiple papers concerned with gene-tree/species-tree reconciliation algorithms (e.g., Vernot et al. 2008; Rasmussen and Kellis 2012; Stolzer et al. 2012). In the presence of ILS, naive reconciliation algorithms place gene duplications on the branch subtending the discordance, with multiple convergent gene losses on descendent branches (Hahn 2007). Newer reconciliation algorithms deal with this bias either by treating the species tree as a polytomy (Vernot et al. 2008; Stolzer et al. 2012) or by assuming there is no hemiplasy (Rasmussen and Kellis 2012). Regardless of the details of these algorithms, it is clear that they offer a way forward for dealing with the mapping of any types of characters onto a “fixed” species trees undergoing ILS (cf. Pollard et al. 2006). Note also that the problem we describe here is not limited to ILS as a cause of discordance: hybridization and introgression between species will also cause discordance, as will errors in phylogenetic reconstruction (Duchêne and Lanfear 2015). All of these will also lead to SPILS-like patterns. However, unlike in the case of ILS – where the proportion of each discordant topology can be predicted (Degnan and Salter 2005) – in the case of introgression one specific discordant topology may be overrepresented. In these situations the set of branches whose lengths are affected is much less predictable.

Finally, it is important to recognize the effect of SPILS on inferences of branch lengths when constructing species trees. Often these branch lengths are used as a first indicator of substitution rate variation, especially if questions concerning the molecular clock are relevant; subsequently they are also important in dating events on a tree. Though the practice of concatenation in species tree inference has received a lot of attention (Edwards et al. 2007; Gatesy and Springer 2014), most of the concern about this approach has involved the accuracy of the tree topology in the “anomaly zone” (Degnan and Rosenberg 2006; Kubatko and Degnan 2007). But concatenation will also force discordant substitutions to be resolved on the most common topology, which will result in SPILS. This problem will be amplified as one concatenates a greater number of loci and a greater number of taxa. Multiple coalescent methods have been developed to overcome the topology-based problems with concatenation (e.g., Drummond and Rambaut 2007; Liu et al. 2009, 2010; Larget et al. 2010; Bryant et al. 2012; Mirarab and Warnow 2015; Yang 2015), but some of these do not estimate branch lengths (e.g., Mirarab and Warnow 2015), and even those that do may report lengths in coalescent units (e.g., Larget et al. 2010; Liu et al. 2010). Moving forward, it is clear that more methods that can simultaneously account for the topological discordance and branch-length discrepancies caused by ILS will be needed, and that they will need to be scalable to genome data. Only with such methods in hand will we be able to get an accurate picture of true substitution rate heterogeneity across large clades.

## Acknowledgements

We thank the members of the Hahn lab for helpful discussions. This work was supported by National Science Foundation grants DBI-0845494 and DEB-1136707.

---

Supplementary Figure Captions

**Supplementary Figure 1.**
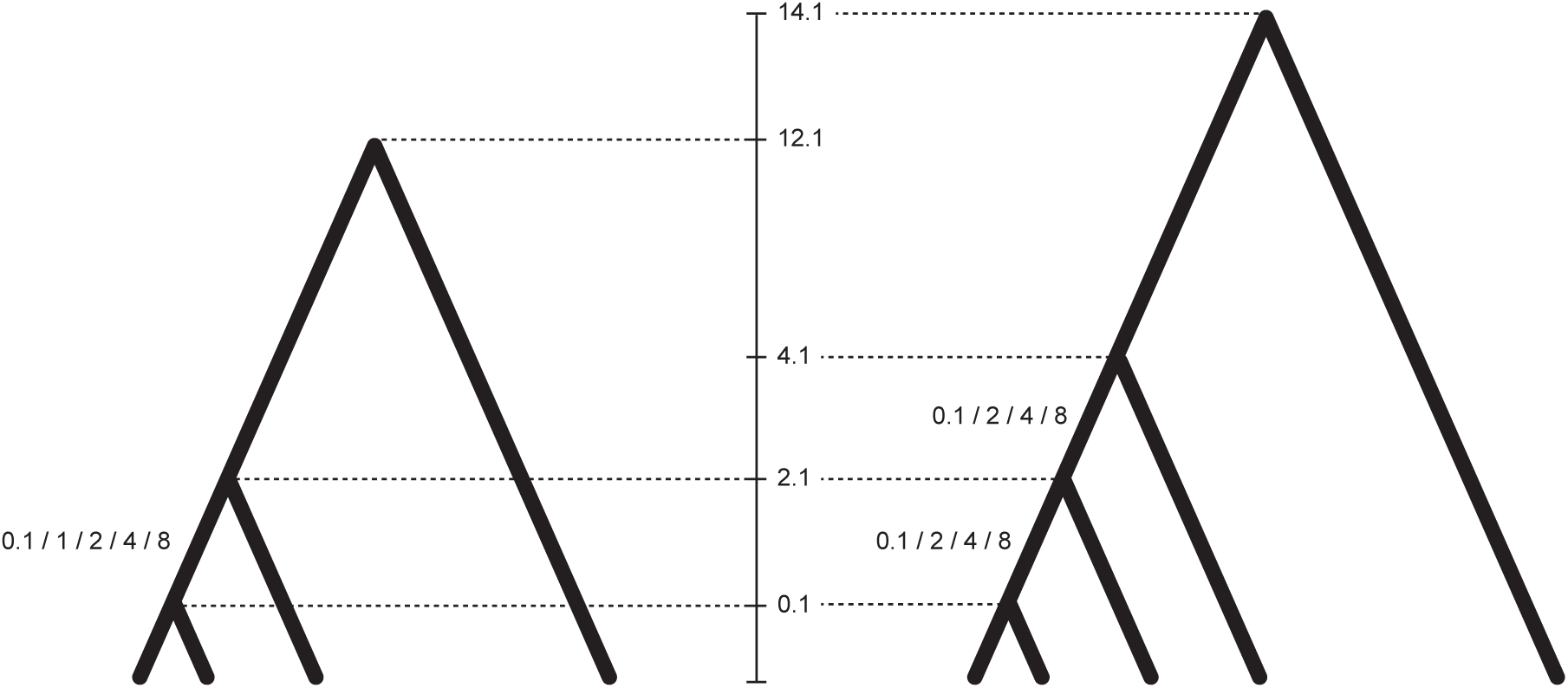
Simulation parameters. The vertical scale shows divergence times in coalescent units. Numbers beside internal branches are factors by which their *N_e_*’s were multiplied in order to produce varying levels of ILS.

**Supplementary Figure 2.**
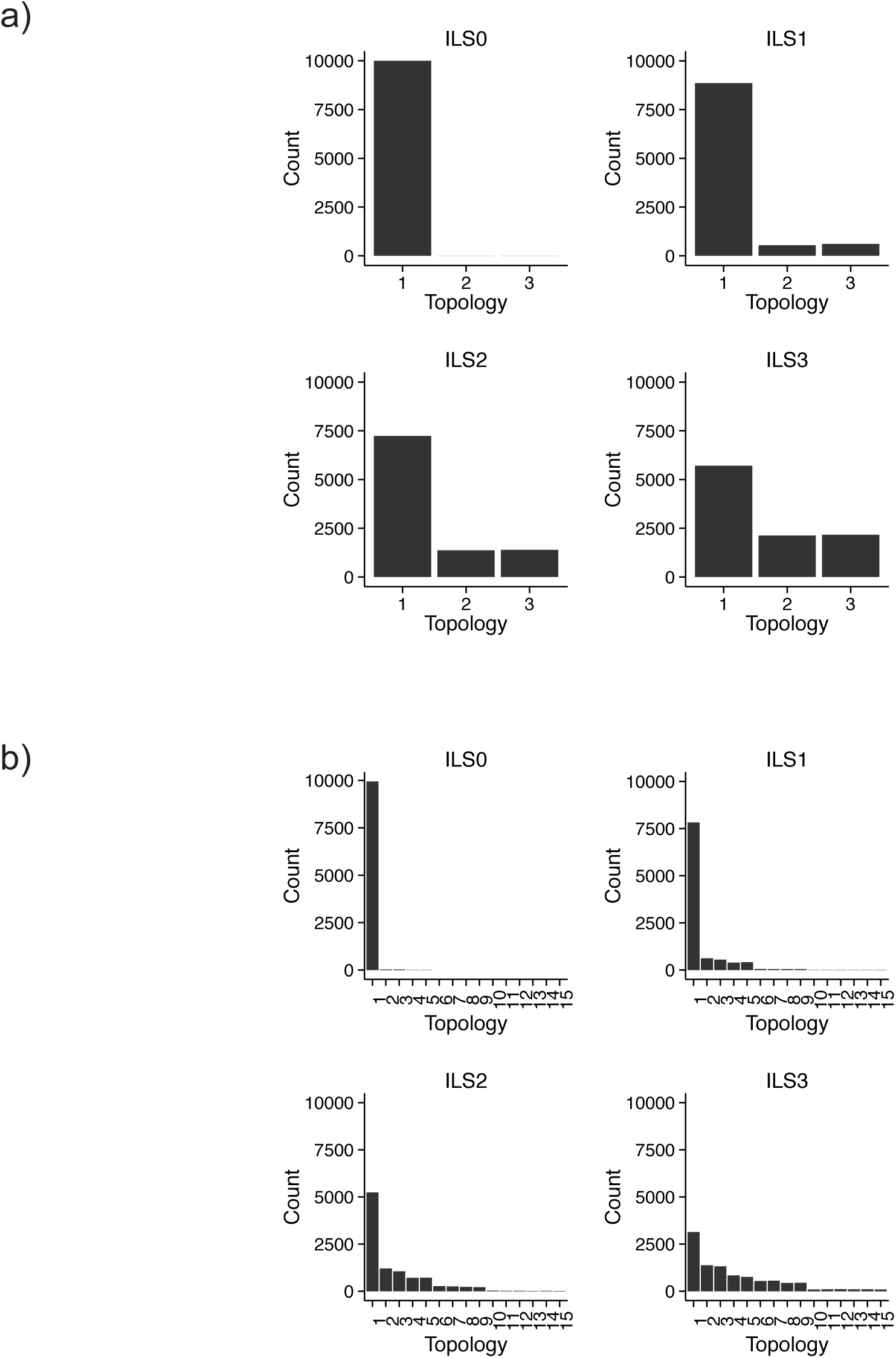
Gene tree topology counts for ILS among a) ((A,B),C), and b) (((A,B),C),D). Topologies are numbered from most to least frequent.

**Supplementary Figure 3.**
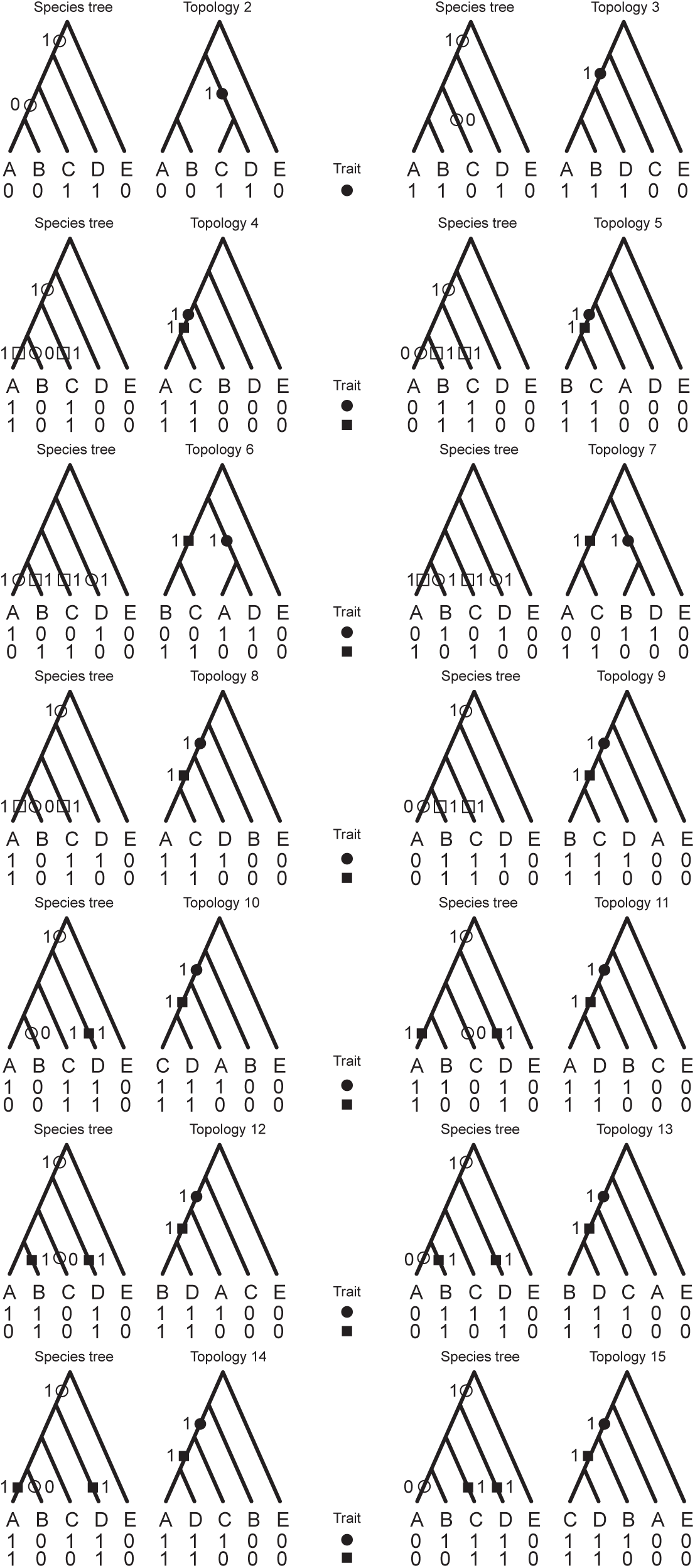
Substitution placement on branches of species tree (left) for each of the 14 possible discordant tree topologies in the presence of ILS (right). Filled symbols correspond to 0➔1 substitutions that occurred on the discordant genealogy of a locus, hollow symbols represent substitutions incorrectly mapped to the species tree.

**Supplementary Figure 4.**
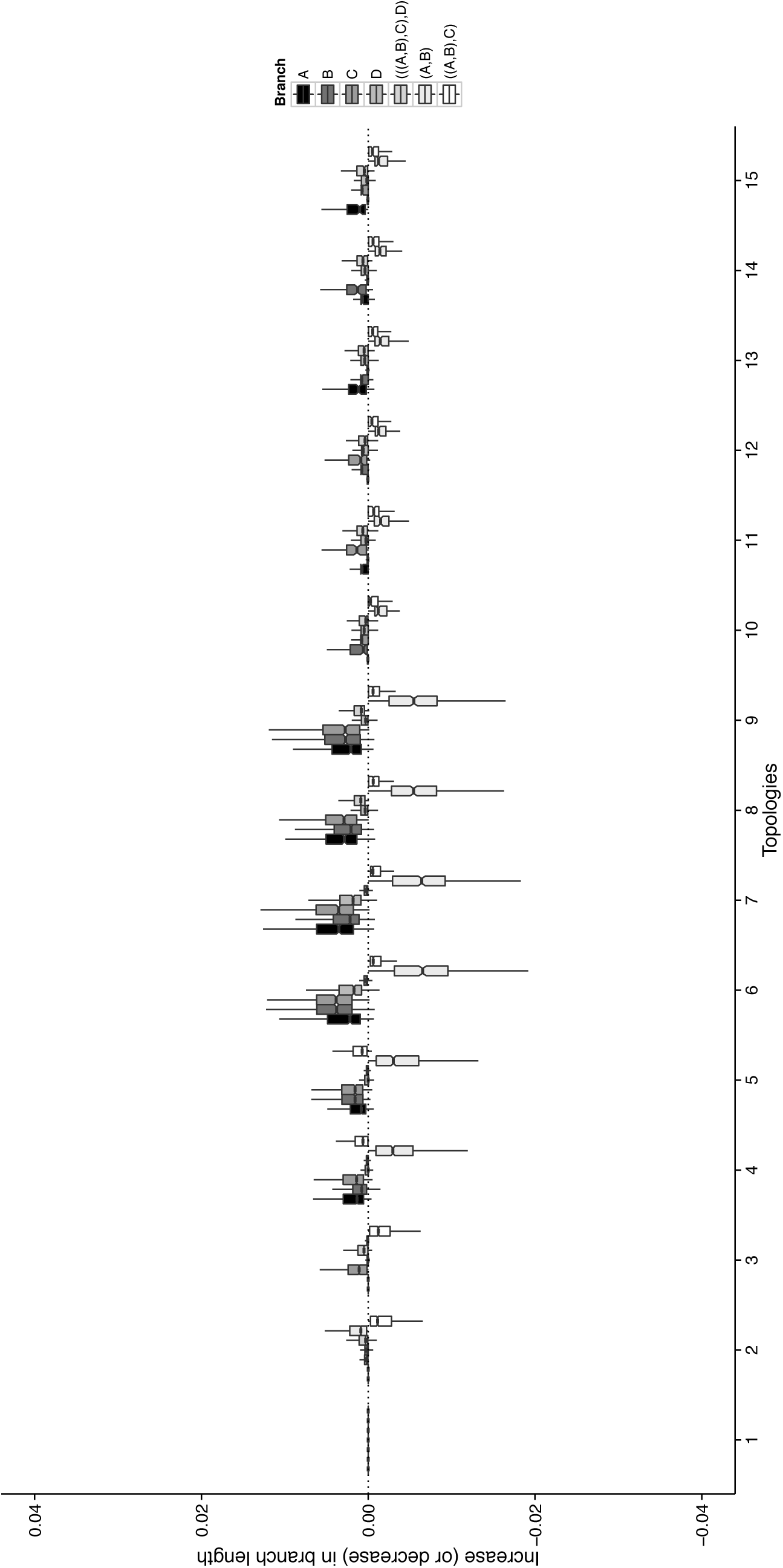
Increase or decrease in branch lengths by topology when ILS is present for species tree ((((A,B),C),D),E). Branch length differences (F – U) come from 10,000 simulated loci.

**Supplementary Figure 5.**
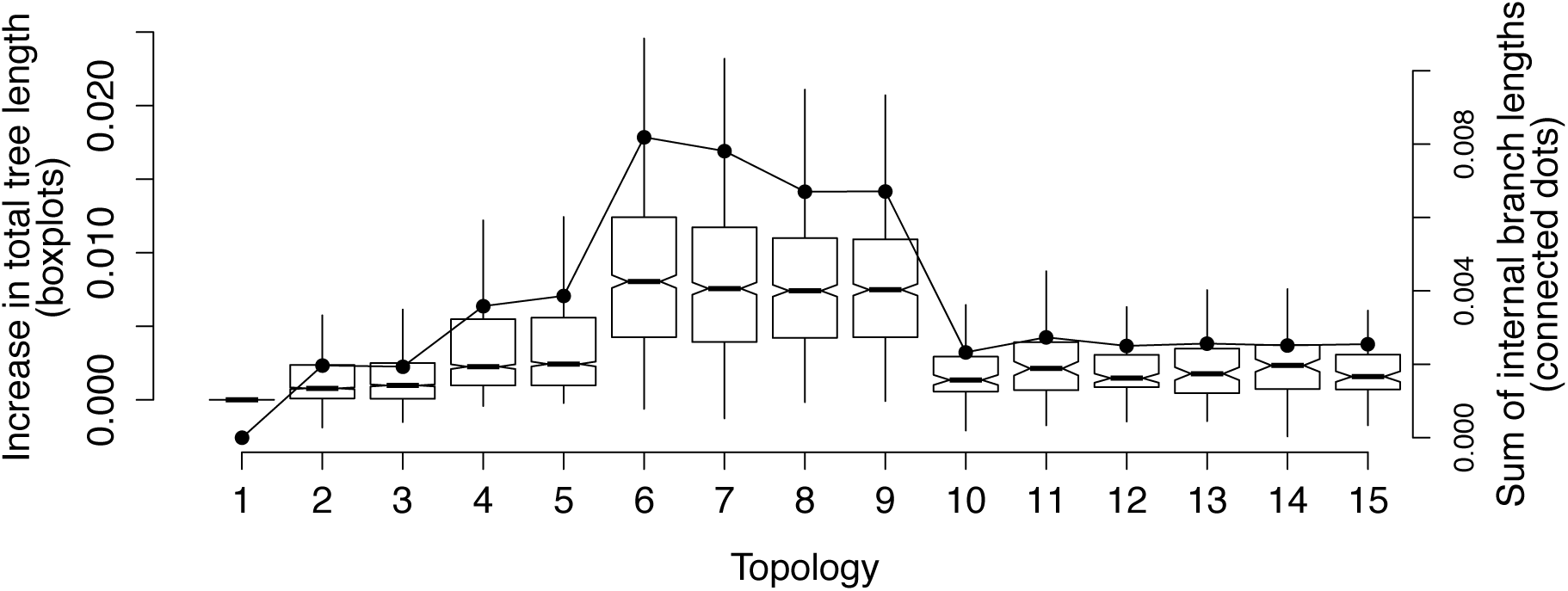
Total tree length increase or decrease (F – U) by tree topology (left axis). Sum of internal branches on each tree topology not present in species tree ((((A,B),C),D),E).

## References

Angelis K., dos Reis M. 2015. The impact of ancestral population size and incomplete lineage sorting on Bayesian estimation of species divergence times. Current Zoology. 61: 1–22.

Avise J., Robinson T. 2008. Hemiplasy: a new term in the lexicon of phylogenetics. Systematic Biology. 57: 503–7.

Barraclough T., Savolainen V. 2001. Evolutionary rates and species diversity in flowering plants. Evolution. 55: 677–83.

Begun D., Holloway A., Stevens K., Hillier L., Poh Y.-P., Hahn M., Nista P., Jones C., Kern A., Dewey C., Pachter L., Myers E., Langley C. 2007. Population Genomics: Whole-Genome Analysis of Polymorphism and Divergence in Drosophila simulans. PLoS Biology. 5: 1–26.

Brawand D., Wagner C., Li Y., Malinsky M., Keller I., Fan S., Simakov O., Ng A., Lim Z., Bezault E., Turner-Maier J., Johnson J., Alcazar R., Noh H., Russell P., Aken B., Alföldi J., Amemiya C., Azzouzi N., Baroiller J.-F., Barloy-Hubler F., Berlin A., Bloomquist R., Carleton K., Conte M., D’Cotta H., Eshel O., Gaffney L., Galibert F., Gante H., Gnerre S., Greuter L., Guyon R., Haddad N., Haerty W., Harris R., Hofmann H., Hourlier T., Hulata G., Jaffe D., Lara M., Lee A., MacCallum I., Mwaiko S., Nikaido M., Nishihara H., Ozouf-Costaz C., Penman D., Przybylski D., Rakotomanga M., Renn S., Ribeiro F., Ron M., Salzburger W., Sanchez-Pulido L., Santos M., Searle S., Sharpe T., Swofford R., Tan F., Williams L., Young S., Yin S., Okada N., Kocher T., Miska E., Lander E., Venkatesh B., Fernald R., Meyer A., Ponting C., Streelman J., Lindblad-Toh K., Seehausen O., Palma F. 2014. The genomic substrate for adaptive radiation in African cichlid fish. Nature. 513: 375–81.

Bromham L. 2009. Why do species vary in their rate of molecular evolution? Biology Letters. 5: 401–4.

Bromham L., Rambaut A., Harvey P.H. 1996. Determinants of rate variation in mammalian DNA sequence evolution. Journal of Molecular Evolution. 43: 610–21.

Bryant D., Bouckaert R., Felsenstein J., Rosenberg N., RoyChoudhury A. 2012. Inferring species trees directly from biallelic genetic markers: bypassing gene trees in a full coalescent analysis. Molecular Biology and Evolution. 29: 1917–32.

Chang BH, Shimmin LC, Shyue SK, Hewett-Emmett D, Li WH. 1994. Weak male-driven molecular evolution in rodents. Proceedings of the National Academy of Sciences of the United States of America. 91: 827–31.

Degnan J., Rosenberg N. 2006. Discordance of species trees with their most likely gene trees. PLoS Genetics. 2: 0762–68.

Degnan J., Salter L. 2005. Gene tree distributions under the coalescent process. Evolution. 63: 24–37.

Drummond A., Rambaut A. 2007. BEAST: Bayesian evolutionary analysis by sampling trees. BMC Evolutionary Biology. 7: 1–8.

Duchene S., Lanfear R. 2015. Phylogenetic uncertainty can bias the number of evolutionary transitions estimated from ancestral state reconstruction methods. Journal of Experimental Zoology Part B: Molecular and Developmental Evolution. 324: 517–24.

Ebersberger I., Metzler D., Schwarz C., Pääbo S. 2002. Genomewide comparison of DNA sequences between humans and chimpanzees. American Journal of Human Genetics. 70: 1490–97.

Edwards S., Liu L., Pearl D. 2007. High-resolution species trees without concatenation. Proceedings of the National Academy of Sciences of the United States of America. 104: 5936–41.

Gatesy J., Springer M. 2014. Phylogenetic analysis at deep timescales: unreliable gene trees, bypassed hidden support, and the coalescence/concatalescence conundrum. Molecular Phylogenetics and Evolution. 80: 231–66.

Gillespie J., Langley C. 1979. Are evolutionary rates really variable? Journal of Molecular Evolution. 13: 27–34.

Good J., Wiebe V., Albert F., Burbano H., Kircher M., Green R., Halbwax M., André C., Atencia R., Fischer A., Pääbo S. 2013. Comparative population genomics of the ejaculate in humans and the great apes. Molecular Biology and Evolution. 30: 964–76.

Guindon S., Gascuel O. 2003. A Simple, Fast, and Accurate Algorithm to Estimate Large Phylogenies by Maximum Likelihood. Systematic Biology. 52: 696704.

Hahn M. 2007. Bias in phylogenetic tree reconciliation methods: implications for vertebrate genome evolution. Genome Biology. 8:R141.

Han M., Thomas G., Lugo-Martinez J., Hahn M. 2013. Estimating gene gain and loss rates in the presence of error in genome assembly and annotation using CAFE 3. Molecular Biology and Evolution. 30: 1987–97.

Hellmann I., Ebersberger I., Ptak S., Pääbo S., Przeworski M. 2003. A neutral explanation for the correlation of diversity with recombination rates in humans. The American Journal of Human Genetics. 72: 1527–35.

Hugall A., Lee M. 2007. The likelihood node density effect and consequences for evolutionary studies of molecular rates. Evolution. 61: 2293–307.

Jarvis E.D., Mirarab S., Aberer A.J., Li B., Houde P., Li C., Ho S.Y., Faircloth B.C., Nabholz B., Howard J.T., Suh A., Weber C.C., da Fonseca R.R., Li J., Zhang F., Li H., Zhou L., Narula N., Liu L., Ganapathy G., Boussau B., Bayzid M.S., Zavidovych V., Subramanian S., Gabaldón T., Capella-Gutiérrez S., Huerta-Cepas J., Rekepalli B., Munch K., Schierup M., Lindow B., Warren W.C., Ray D., Green R.E., Bruford M.W., Zhan X., Dixon A., Li S., Li N., Huang Y., Derryberry E.P., Bertelsen M.F., Sheldon F.H., Brumfield R.T., Mello C.V., Lovell P.V., Wirthlin M., Schneider M.P., Prosdocimi F., Samaniego J.A.A., Vargas Velazquez A.M., Alfaro-Núñez A., Campos P.F., Petersen B., Sicheritz-Ponten T., Pas A., Bailey T., Scofield P., Bunce M., Lambert D.M., Zhou Q., Perelman P., Driskell A.C., Shapiro B., Xiong Z., Zeng Y., Liu S., Li Z., Liu B., Wu K., Xiao J., Yinqi X., Zheng Q., Zhang Y., Yang H., Wang J., Smeds L., Rheindt F.E., Braun M., Fjeldsa J., Orlando L., Barker F.K., Jønsson K.A., Johnson W., Koepfli K.-P.P., O’Brien S., Haussler D., Ryder O.A., Rahbek C., Willerslev E., Graves G.R., Glenn T.C., McCormack J., Burt D., Ellegren H., Alström P., Edwards S.V., Stamatakis A., Mindell D.P., Cracraft J. 2014. Whole-genome analyses resolve early branches in the tree of life of modern birds. Science. 346: 1320–31.

Jónsson H., Schubert M., Seguin-Orlando A., Ginolhac A., Petersen L., Fumagalli M., Albrechtsen A., Petersen B., Korneliussen T., Vilstrup J., Lear T., Myka J., Lundquist J., Miller D., Alfarhan A., Alquraishi S., Al-Rasheid K., Stagegaard J., Strauss G., Bertelsen M., Sicheritz-Ponten T., Antczak D., Bailey E., Nielsen R., Willerslev E., Orlando L. 2014. Speciation with gene flow in equids despite extensive chromosomal plasticity. Proceedings of the National Academy of Sciences of the United States of America. 111: 18655–60.

Kubatko L., Degnan J. 2007. Inconsistency of phylogenetic estimates from concatenated data under coalescence. Systematic Biology. 56: 17–24.

Laird C.D., McConaughy B.L., McCarthy B.J. 1969. Rate of fixation of nucleotide substitutions in evolution. Nature. 224: 149–54.

Lamichhaney S., Berglund J., Almén M., Maqbool K., Grabherr M., Martinez-Barrio A., Promerová M., Rubin C.-J., Wang C., Zamani N., Grant B., Grant P., Webster M., Andersson L. 2015. Evolution of Darwin’s finches and their beaks revealed by genome sequencing. Nature. 518: 371–75.

Lanfear R., Kokko H., Eyre-Walker A. 2014. Population size and the rate of evolution. Trends in Ecology and Evolution. 29: 33–41.

Lanfear R., Welch J.J., Bromham L. 2010. Watching the clock: studying variation in rates of molecular evolution between species. Trends in Ecology and Evolution. 25: 495–503.

Larget B., Kotha S., Dewey C., Ané C. 2010. BUCKy: Gene tree/species tree reconciliation with Bayesian concordance analysis. Bioinformatics. 26: 2910–11.

Larracuente A., Sackton T., Greenberg A., Wong A., Singh N., Sturgill D., Zhang Y., Oliver B., Clark A. 2008. Evolution of protein-coding genes in Drosophila. Trends in Genetics. 24: 114–23.

Li W.-H. 1997. Molecular evolution. Sinauer Associates.

Liu L., Yu L., Edwards S. 2010. A maximum pseudo-likelihood approach for estimating species trees under the coalescent model. BMC Evolutionary Biology. 10: 1–18.

Liu L., Yu L., Pearl D., Edwards S. 2009. Estimating species phylogenies using coalescence times among sequences. Systematic Biology. 58: 468–77.

Lynch M. 2010. Evolution of the mutation rate. Trends in Genetics. 26: 345–52.

Maddison W.P. 1997. Gene trees in species trees. Systematic Biology. 46: 523–36.

Makova K.D., Li W.-H.H. 2002. Strong male-driven evolution of DNA sequences in humans and apes. Nature. 416: 624–6.

Malcom C.M., Wyckoff G.J., Lahn B.T. 2003. Genic mutation rates in mammals: local similarity, chromosomal heterogeneity, and X-versus-autosome disparity. Molecular Biology and Evolution. 20: 1633–41.

Martin A., Palumbi S. 1993. Body size, metabolic rate, generation time, and the molecular clock. Proceedings of the National Academy of Sciences of the United States of America. 90: 4087–91.

Mirarab S., Warnow T. 2015. ASTRAL-II: coalescent-based species tree estimation with many hundreds of taxa and thousands of genes. Bioinformatics. 31:i44–i52.

Mita S., Siol M. 2012. EggLib: processing, analysis and simulation tools for population genetics and genomics. BMC Genetics. 13: 1–12.

Miyata T., Hayashida H., Kuma K., Mitsuyasu K., Yasunaga T. 1987. Male-driven molecular evolution: a model and nucleotide sequence analysis. Cold Spring Harbor symposia on quantitative biology. 52: 863–67.

Oliver J.C. 2013. Microevolutionary processes generate phylogenomic discordance at ancient divergences. Evolution. 67: 1823–30.

Pagel M., Venditti C., Meade A. 2006. Large punctuational contribution of speciation to evolutionary divergence at the molecular level. Science. 314: 119–21.

Pamilo P., Nei M. 1988. Relationships between gene trees and species trees. Molecular Biology and Evolution. 5: 568–83.

Parker J., Tsagkogeorga G., Cotton J., Liu Y., Provero P., Stupka E., Rossiter S. 2013. Genome-wide signatures of convergent evolution in echolocating mammals. Nature. 502: 228–31.

Pease J., Hahn M. 2013. More accurate phylogenies inferred from low-recombination regions in the presence of incomplete lineage sorting. Evolution. 67: 2376–84.

Pollard D.A., Iyer V.N., Moses A.M., Eisen M.B. 2006. Widespread discordance of gene trees with species tree in Drosophila: evidence for incomplete lineage sorting. PLoS Genetics. 2: 1634–47.

Rasmussen M., Kellis M. 2012. Unified modeling of gene duplication, loss, and coalescence using a locus tree. Genome Research. 22: 755–65.

Robinson D.F., Foulds L.R. 1981. Comparison of phylogenetic trees. Mathematical Biosciences. 53: 131–47.

Scally A., Dutheil J., Hillier L., Jordan G., Goodhead I., Herrero J., Hobolth A., Lappalainen T., Mailund T., Marques-Bonet T., McCarthy S., Montgomery S., Schwalie P., Tang Y., Ward M., Xue Y., Yngvadottir B., Alkan C., Andersen L., Ayub Q., Ball E., Beal K., Bradley B., Chen Y., Clee C., Fitzgerald S., Graves T., Gu Y., Heath P., Heger A., Karakoc E., Kolb-Kokocinski A., Laird G., Lunter G., Meader S., Mort M., Mullikin J., Munch K., O’Connor T., Phillips A., Prado-Martinez J., Rogers A., Sajjadian S., Schmidt D., Shaw K., Simpson J., Stenson P., Turner D., Vigilant L., Vilella A., whitener W., Zhu B., Cooper D., Jong P., Dermitzakis E., Eichler E., Flicek P., Goldman N., Mundy N., Ning Z., Odom D., Ponting C., Quail M., Ryder O., Searle S., Warren W., Wilson R., Schierup M., Rogers J., Tyler-Smith C., Durbin R. 2012. Insights into hominid evolution from the gorilla genome sequence. Nature. 483: 169–75.

Shimmin L.C., Chang B.H.-J., Li W.-H. 1993. Male-driven evolution of DNA sequences. Nature. 362: 745–47.

Smith S., Donoghue M. 2008. Rates of molecular evolution are linked to life history in flowering plants. Science. 322: 86–89.

Stamatakis A. 2014. RAxML version 8: a tool for phylogenetic analysis and post-analysis of large phylogenies. Bioinformatics. 30: 1312–13.

Stolzer M., Lai H., Xu M., Sathaye D., Vernot B., Durand D. 2012. Inferring duplications, losses, transfers and incomplete lineage sorting with nonbinary species trees. Bioinformatics. 28:i409–15.

Studer R., Penel S., Duret L., Robinson-Rechavi M. 2008. Pervasive positive selection on duplicated and nonduplicated vertebrate protein coding genes. Genome Research. 18: 1393–1402.

Suh A., Smeds L., Ellegren H. 2015. The dynamics of incomplete lineage sorting across the ancient adaptive radiation of neoavian birds. PLoS Biology. 13: 1–18.

The Chimpanzee Sequencing and Analysis Consortium. 2005. Initial sequence of the chimpanzee genome and comparison with the human genome. Nature. 437: 69–87.

Venditti C., Pagel M. 2009. Speciation as an active force in promoting genetic evolution. Trends in Ecology and Evolution. 25: 14–20.

Vernot B., Stolzer M., Goldman A., Durand D. 2008. Reconciliation with non-ninary species trees. Journal of Computational Biology. 15: 981–1006.

Webster A., Payne R., Pagel M. 2003. Molecular phylogenies link rates of evolution and speciation. Science. 301: 478.

Wilson Sayres M.A., Venditti C., Pagel M., Makova K.D. 2011. Do variations in substitution rates and male mutation bias correlate with life-history traits? A study of 32 mammalian genomes. Evolution. 65: 2800–15.

Witt C., Brumfield R. 2004. Comment on “Molecular Phylogenies Link Rates of Evolution and Speciation”. Science. 303:173b.

Wright S., Keeling J., Gillman L. 2006. The road from Santa Rosalia: a faster tempo of evolution in tropical climates. Proceedings of the National Academy of Sciences of the United States of America. 103: 7718–22.

Wu C., Li W. 1985. Evidence for higher rates of nucleotide substitution in rodents than in man. Proceedings of the National Academy of Sciences of the United States of America. 82: 174–145.

Yang Z. 2007. PAML 4: Phylogenetic Analysis by Maximum Likelihood. Molecular Biology and Evolution. 24: 1586–91.

Yang Z. 2015. The BPP program for species tree estimation and species delimitation. Current Zoology. 61: 854–65.

Zhang G., Li C., Li Q., Li B., Larkin D., Lee C., Storz J., Antunes A., Greenwold M., Meredith R., Ödeen A., Cui J., Zhou Q., Xu L., Pan H., Wang Z., Jin L., Zhang P., Hu H., Yang W., Hu J., Xiao J., Yang Z., Liu Y., Xie Q., Yu H., Lian J., Wen P., Zhang F., Li H., Zeng Y., Xiong Z., Liu S., Zhou L., Huang Z., An N., Wang J., Zheng Q., Xiong Y., Wang G., Wang B., Wang J., Fan Y., Fonseca R., Alfaro-Núñez A., Schubert M., Orlando L., Mourier T., Howard J., Ganapathy G., Pfenning A., Whitney O., Rivas M., Hara E., Smith J., Farré M., Narayan J., Slavov G., Romanov M., Borges R., Machado J., Khan I., Springer M., Gatesy J., Hoffmann F., Opazo J., Håstad O., Sawyer R., Kim H., Kim K.-W., Kim H., Cho S., Li N., Huang Y., Bruford M., Zhan X., Dixon A., Bertelsen M., Derryberry E., Warren W., Wilson R., Li S., Ray D., Green R., O’Brien S., Griffin D., Johnson W., Haussler D., Ryder O., Willerslev E., Graves G., Alström P., Fjeldså J., Mindell D., Edwards S., Braun E., Rahbek C., Burt D., Houde P., Zhang Y., Yang H., Wang J., Jarvis E., Gilbert M., Wang J., Ye C., Liang S., Yan Z., Zepeda M., Campos P., Velazquez A., Samaniego J., Avila-Arcos M., Martin M., Barnett R., Ribeiro A., Mello C., Lovell P., Almeida D., Maldonado E., Pereira J., Sunagar K., Philip S., Dominguez-Bello M., Bunce M., Lambert D., Brumfield R., Sheldon F., Holmes E., Gardner P., Steeves T., Stadler P., Burge S., Lyons E., Smith J., McCarthy F., Pitel F., Rhoads D., Froman D. 2014. Comparative genomics reveals insights into avian genome evolution and adaptation. Science. 346: 1311–20.

